# Multiomic single-cell sequencing defines tissue-specific responses in Stevens-Johnson Syndrome and Toxic epidermal necrolysis

**DOI:** 10.1101/2023.11.26.568771

**Authors:** Andrew Gibson, Ramesh Ram, Rama Gangula, Yueran Li, Eric Mukherjee, Amy M Palubinsky, Chelsea N Campbell, Michael Thorne, Katherine C Konvinse, Phuti Choshi, Pooja Deshpande, Sarah Pedretti, Richard T O’Neil, Celestine N Wanjalla, Spyros A Kalams, Silvana Gaudieri, Rannakoe J Lehloenya, Samuel S Bailin, Abha Chopra, Jason A Trubiano, AUS-SCAR study group, Jonny G Peter, AFRI-SCAR and IMARI-Africa study group, Simon A Mallal, Elizabeth J Phillips

## Abstract

Stevens-Johnson syndrome and toxic epidermal necrolysis (SJS/TEN) is a rare but life-threatening cutaneous drug reaction mediated by human leukocyte antigen (HLA) class I-restricted CD8+ T-cells. To obtain an unbiased assessment of SJS/TEN cellular immunopathogenesis, we performed single-cell (sc) transcriptome, surface proteome, and TCR sequencing on unaffected skin, affected skin, and blister fluid from 17 SJS/TEN patients. From 119,784 total cells, we identified 16 scRNA-defined subsets, confirmed by subset-defining surface protein expression. Keratinocytes upregulated HLA and IFN-response genes in the affected skin. Cytotoxic CD8+ T-cell subpopulations of expanded and unexpanded TCRαβ clonotypes were shared in affected skin and blister fluid but absent or unexpanded in SJS/TEN unaffected skin. SJS/TEN blister fluid is a rich reservoir of oligoclonal CD8+ T-cells with an effector phenotype driving SJS/TEN pathogenesis. This multiomic database will act as the basis to define antigen-reactivity, HLA restriction, and signatures of drug-antigen-reactive T-cell clonotypes at a tissue level.

## Main

Stevens-Johnson syndrome and toxic epidermal necrolysis (SJS/TEN) is a rare T-cell-mediated severe cutaneous adverse drug reaction characterized by skin and mucous membrane blistering and detachment. The hallmarks of SJS/TEN are keratinocyte death and mortality up to 50%^1^. Human leukocyte antigen (HLA) class I alleles are strongly associated with drug-induced SJS/TEN^2^ and drug-antigen-expanded T-cell receptor (TCR) clonotypes are identified in patient blister fluid that are rare in the peripheral blood and expressed on CD8+ T-cells producing granulysin^3^; thought to be the primary mediator of keratinocyte death^4^. However, the signatures of antigen-driven TCR-expressing T-cell population(s) at the site of tissue damage, which will elucidate tissue-relevant biological markers for earlier diagnosis and treatment, cannot be determined without unbiased single-cell analyses of normal and diseased skin. To define the immunopathogenesis of SJS/TEN and tissue-relevant signatures of oligoclonal drug-antigen-reactive T-cells, we performed single-cell transcriptome (scRNA-seq), surface protein (scCITE-seq), and TCR-sequencing across unaffected skin, affected skin, and blister fluid from 17 patients with SJS/TEN. Patients varied in age, sex, culprit drug, and treatment. Normal skin (n=1) and burn blister fluid (n=1) were included as controls.

The total scRNA-defined uniform manifold approximation and projection (UMAP, Fig. 1Ai) includes 119,784 cells, which span 16 subtypes, including CD8+ and CD4+ T-cells, natural killer (NK) cells, B-cells, monocytes, macrophages, dendritic cells (DC), and keratinocytes (Fig. 1Aii). ScCITE-seq confirmed unsupervised transcriptome-based clustering with characteristic protein expression across scRNA-defined CD3+ CD4+ and CD8+ T-cells, CD56+ NK cells, CD1c+ DC, CD19+ B-cells, and CD14+ CD11b+ monocytes and macrophages (Fig. 1Bi-ii). Surface phenotyping identified that few NK or T-cells expressed CD3 or CD56, respectively, suggesting an absence of NKT-cells, and that myeloid subsets in SJS/TEN were pro-inflammatory, including DC (CD86+, CD112+) but also monocytes and macrophages (CD40+, CD54+, CD31+; Extended Data Fig. 1). Macrophages expressed a mixed pro-inflammatory M1-(CD11c+) and anti-inflammatory M2a/M2c (CD11b+, CD163+)-like surface phenotype. Transcriptomic analyses confirmed an intermediate-like macrophage signature (*CD9*^*+*^, *CD163*^*+*^ *APOE*^*+*^ *APOC1*^*+*^)^5^, supporting a mixed surface phenotype and subpopulations involved in inflammation and others with repair.

**Figure 1.**
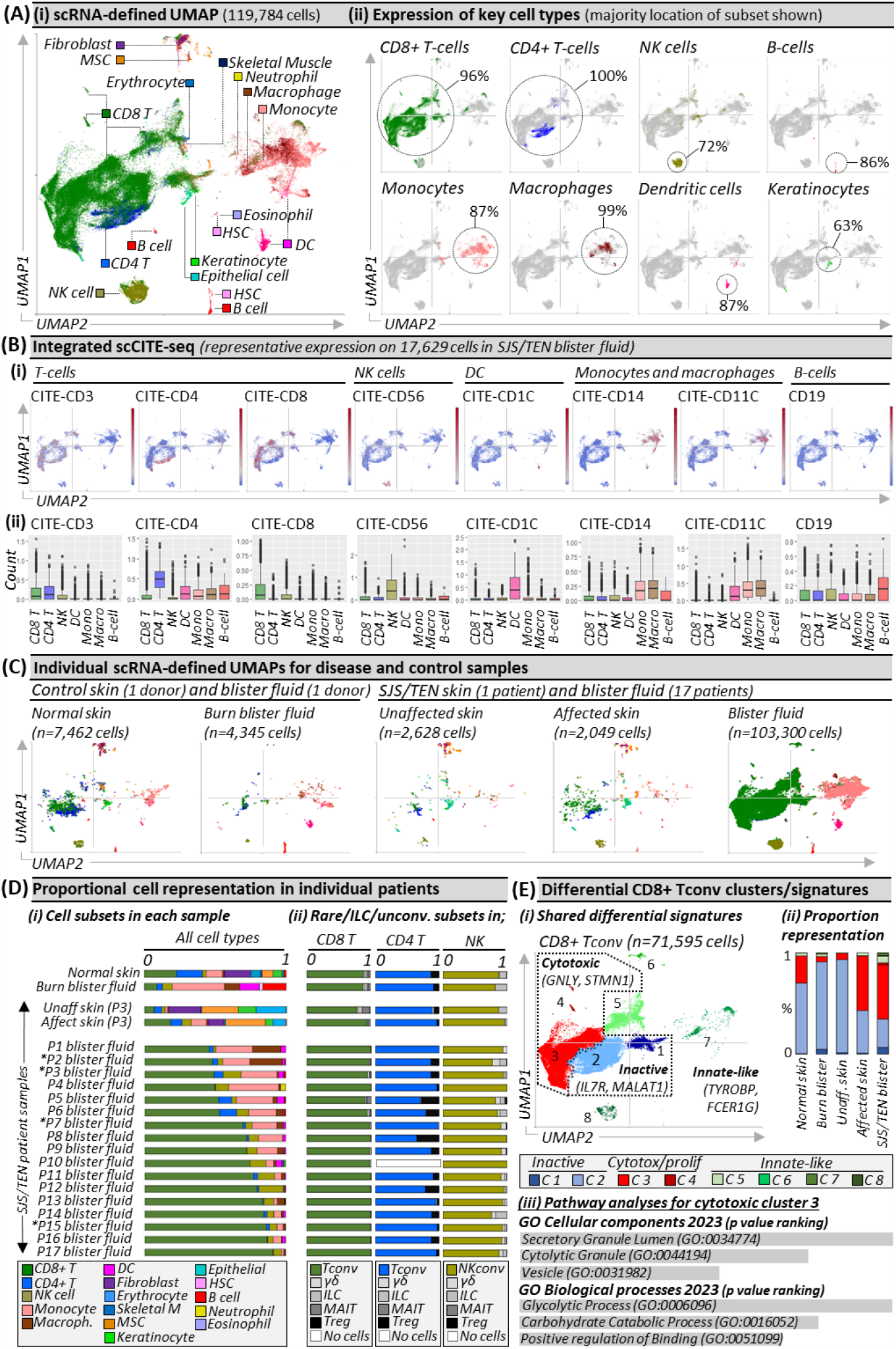
Unbiased multi-focal single-cell sequencing identifies expanded pathogenic CD8+ Tconv in affected skin and blister fluid during Stevens-Johnson syndrome and toxic epidermal necrolysis. (A) (i) scRNA-defined uniform manifold approximation and projection (UMAP) of 119,784 live cells from normal skin, burn blister fluid, and unaffected skin, affected skin and/or blister fluid from 17 patients with acute SJS/TEN. (ii) Expression distribution for key cell types. The majority UMAP location of each subset is labeled with the percentage of each subset within that location indicated. Figure created using VGAS. (B) Integrated scCITE-seq expression of subset-defining cell surface proteins for T-cells (CD3, CD4, CD8), NK cells (CD56), DC (CD1c), monocytes and macrophages (CD14, CD11c), and B cells (CD19). (i) Heatmap expression (high to low, red to blue) for each surface protein is shown on a representative UMAP of 17,629 cells obtained within a single 10x run, and (ii) average box plot expression is shown between scRNA-defined subsets. Figure created using VGAS. (C) Individual scRNA-defined UMAPs for tissue-relevant controls from unrelated donors (burn blister fluid, normal skin) and skin and blister fluid from patients with SJS/TEN. The average expression is shown for patients with multiple samples from the same time point, indicated by an asterisk. Figure created using VGAS. (D) scRNA-defined cell representation of (i) all subsets in individual samples and (ii) rare, unconventional, and innate-like lymphoid cell (ILC) populations as a proportion of total CD8 T-cells, CD4 T-cells, or NK cells in each sample. (E) (i) Seurat-defined Tconv clusters 1-8 with shared naming according to differential gene expression signatures and (ii) proportional representation in control and disease sample phenotypes. (iii) Pathway analysis for the significant differentially expressed gene signature of CD8 Tconv cluster 3 using Enrichr and gene ontology (GO) cellular and biological terms ranked by p-value. *MSC, Mesenchymal stromal cell; HSC, Hematopoietic stem cell; NK, Natural killer; DC, Dendritic cell; Tconv, T conventional cell; unconv*., *unconventional; ILC, innate-like lymphoid cell; Unaff, Unaffected; Aff, Affected*.

To determine subsets enriched in disease compared to control phenotypes (Fig. 1C), we compared proportional representation. The most prevalent immune population in SJS/TEN-affected skin and blister fluid were CD8+ T-cells (Fig. 1Di), which were the only significantly enriched subset in SJS/TEN blister fluid (n=17) compared to grouped controls (burn blister fluid, n=1; normal skin, n=1; unaffected SJS/TEN patient skin, n=1) (scCODA, p<0.05). On average, SJS/TEN blister fluid contained 69% CD8+ T-cells, 14% monocytes, 8% NK cells, 4% macrophages, 4% CD4+ T-cells, and 1% DC. CD8+ T-cells (18%) were also the most abundant immune subset in affected skin, with a 2.7-fold increase in CD8+ T-cells in affected compared to unaffected skin (Fig. 1Di). Consistent with previous studies, B-cells represented <0.5% of cells in SJS/TEN-affected skin and blister fluid^6,7^.

While conventional T-cells recognizing protein-derived antigens (Tconv) are implicated as the prevalent cytotoxic subset in SJS/TEN skin^7^, rare, unconventional, and innate-like lymphoid cell (ILC) populations are important immuno-inflammatory regulators but may not always be detected^8^. To elucidate effector phenotype, we realigned T-and NK cells to the Azimuth peripheral blood mononuclear cell (PBMC) reference^9^, which contains T regulatory (Treg) cells, ILC, and unconventional subsets including gamma delta (γδ) T-cells and mucosal-associated invariant (MAI)T-cells. Although most cells did not re-align with rare, ILC, or unconventional subsets, small populations were identified (Fig1. Dii). The largest of these were Tregs aligned to the CD4+ T-cell subset in SJS/TEN blister fluid (16%), unaffected skin (8%), and affected skin (4%). Further, ILC comprised >10% of the NK subset in normal skin and burn blister fluid, but <5% of those in SJS/TEN-affected skin and blister fluid. In the broadly-defined CD8+ T-cell subset, ILC and unconventional populations each represented ≤1% of cells in SJS/TEN blister fluid. Overall, 94% and 98% of CD8+ T-cells in SJS/TEN-affected skin and blister fluid were defined as Tconv (Fig1. Dii).

Within the majority CD8+ Tconv population, eight Seurat-defined clusters (Fig. 1Ei) shared differential signatures (Extended Data Table 1) associated with quiescence/inactivation (clusters 1-2; *MALAT1 and/or IL7R*)^10^, cytotoxicity and proliferation (clusters 3-4; *GNLY, STMN1*), and innate-like differentiation (clusters 5-8; *TYROBP, FCER1G*)^11^. Notably, ‘innate-like’ clusters were represented by few cells and aligned to UMAP positions associated with other subsets, suggestive of physically-interacting cells (PIC)^12^. ‘Inactive’ cluster 1 showed high expression of mitochondrial genes without cytotoxic activation, suggesting cell death through basal turnover, and represented 1% of CD8+ Tconv in normal, affected, and unaffected skin. ‘Inactive’ cluster 2 had high expression of *IL7R*, which is downregulated upon antigen-induced activation, indicating a population unstimulated by the drug-antigen during SJS/TEN, and was the predominant CD8+ Tconv subset in normal skin, unaffected skin, and burn blister fluid (Fig. 1Eii). In contrast, ‘cytotoxic’ cluster 3 was the predominant subset in SJS/TEN-affected skin and blister fluid. Pathway analyses using differentially expressed genes (DEG) identified processes including cytotoxic secretion (Fig. 1Eiii), aligning CD8+ Tconv cluster 3 with the immunopathogenesis of SJS/TEN.

Clonally-expanded HLA-class I-restricted TCRαβ CD8+ T-cells are implicated as the drug antigen-driven effectors of SJS/TEN^3,13^. To define signatures of clonally-expanded CD8+ Tconv and whether those in blister fluid represent those in the affected skin, we assessed time-paired samples from a single untreated patient (Fig. 2), including unaffected and affected skin and three blister fluid samples from the arm, face, and foot (Fig. 2Ai). Initial analyses of total CD8+ Tconv in affected compared to unaffected skin identified significant DEG (Fig. 2Aii, Extended Data Table 2) associated with cytotoxicity (*GNLY*), activation/differentiation (*CD27*), regulation (*LAG3*), and the immunoproteasome (*PSMB9*). In contrast, in keratinocytes, upregulated genes were associated with stress-induced migration (*GSTP1, KRT6B*), IFN-induced response (*IFITM3, IFI6*), and HLA class I-associated (*HLA-B, HLA-C*) proteasomal activation (*PSME1, PSME2, PSMB9*), suggesting increased capability to present CD8+ T-cell-directed epitopes (Fig. 2Aii, Extended Data Table 2).

**Figure 2.**
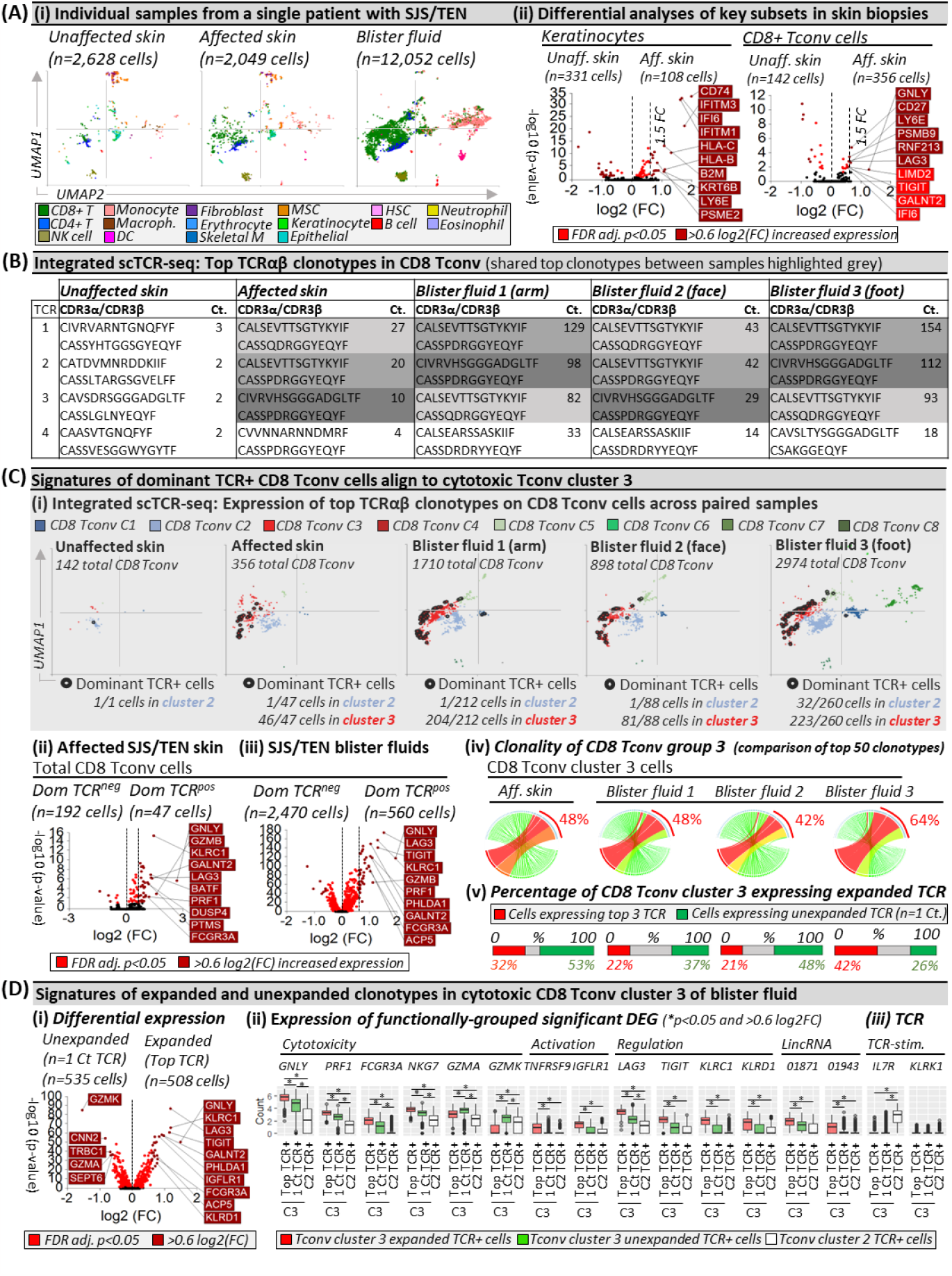
Shared oligoclonal TCRαβ clonotypes on cytotoxic CD8 Tconv in affected skin and blister fluid are absent from or unexpanded in unaffected skin. (A) (i) UMAPs for paired unaffected and affected skin and blister fluid samples from a single patient with SJS/TEN. (ii) Differential gene expression (Wilcoxon, Hochberg adj, p<0.05) between keratinocytes or CD8+ T-cells from unaffected and affected skin. Genes colored red are significantly (p<0.05) increased (light red <0.6log2FC, dark red >0.6log2FC). The top 10 genes are labeled. (B) Top functional TCR CDR3αβ pairings and counts (Ct.) in CD8 Tconv cells. The same top TCR clonotypes across affected samples are highlighted grey. (C) Signatures of dominant TCR+ CD8 Tconv align to cytotoxic Tconv cluster 3. (i) Expression of the top 3 dominantly-expanded TCR (black) on CD8 Tconv cells of each sample. The number of dominant TCR+ cells aligned to CD8 Tconv cluster 2 (control) and cluster 3 (cytotoxic) are indicated numerically. (ii-iii) Differential gene expression (Wilcoxon, Hochberg adj, p<0.05) between CD8 Tconv expressing dominant TCRs or other TCRs in (ii) affected skin or (iii) blister fluids. Genes colored red are significantly (p<0.05) increased (light red <0.6log2FC, dark red >0.6log2FC). The top 10 genes are labeled. (iv) Clonality of CD8 Tconv cluster 3 in affected skin and blister fluid samples. The top 50 TCRs are shown and the percentage indicates the proportion of counts aligned to the top 3 dominantly-expanded TCRs. Circos plot segment width is proportionate to dominance (increasing green to red). (v) Proportional expression of dominantly-expanded TCR+ cells (red) and unexpanded clonotypes (green, n=1 count) in CD8+ Tconv cluster 3 across affected skin and blister fluid samples. (D) Signatures of expanded and unexpanded clonotypes in Tconv cluster 3. (i) Differential gene expression (Wilcoxon, Hochberg adj, p<0.05) between expanded or unexpanded TCR+ cells of CD8 Tconv cluster 3 in SJS/TEN blister fluid. Genes colored red are significantly (p<0.05) increased (light red <0.6log2FC, dark red >0.6log2FC). The top 10 genes are labeled. (ii) DEG and (iii) interest TCR-activation-related gene expression between cells of Tconv cluster 3 expressing expanded (red) or unexpanded (green) TCR compared to TCR from control Tconv cluster 2 (white). *Indicates significant differential expression (Wilcoxon, Hochberg adj, p<0.05 and >0.6 log2FC). Figure created using VGAS. *TCR, T-cell receptor; Dom TCR, Dominantly-expanded TCR; MSC, Mesenchymal stromal cell; HSC, Hematopoietic stem cell; NK, Natural killer; DC, Dendritic cell; Tconv, T conventional cell; CDR3, Complementary determining region 3; Ct*., *count; FDR, false discovery rate; FC, fold change; LincRNA, long intergenic non-coding RNA; Unaff, Unaffected; Aff, Affected*.

Across paired samples from this patient, a productive TCRαβ pair was identified for 67% and 53% of CD4+ and CD8+ T-cells, respectively. The most highly expressed TCRαβ variable genes across CD8+ Tconv of affected samples were *TRAV19* and *TRBV27* (Extended Data Fig. 2) and analyses of paired complementary-determining region (CDR)3αβ identified the same top three dominantly-expanded TCRs across affected skin and blister fluids (Fig. 2B, Extended Data Fig. 3). These clonotypes were not expanded in unaffected skin (Fig. 2B) nor expressed by CD4+ Tconv cells, and the identification of oligoclonal clonotypes in this patient was representative of CD8 Tconv in blister fluid from all SJS/TEN patients (Extended Data Fig. 4). Interestingly, in this patient, the three dominant clonotypes had similar CDR3 sequences, with two sharing a CDR3β (CASSPDRGGYEQYF) but different CDR3α and absent from unaffected skin. A third, expressed on one cell in unaffected skin (Extended Data Fig. 3), had one CDR3β mismatch (CASS<u>Q</u>DRGGYEQYF). GLIPH2 predicted that all three TCRs share peptide binding specificity (p=7.90E-15). Notably, these three dominantly-expanded clonotypes were expressed in the same UMAP location, with the majority of those in affected skin (>97%) and blister fluid (>90%) aligned to cytotoxic CD8+ Tconv cluster 3 (Fig. 2Ci). Compared to CD8+ Tconv cells expressing all other TCR, the same DEG signature (Extended Data Table 3) of dominantly-expanded TCR+ CD8+ Tconv in affected skin (Fig. 2Cii) and blister fluid (Fig. 2Ciii) aligned with mature cytotoxic effectors (*GNLY, GZMB, PRF1, FCGR3A*), cytolytic granule exocytosis (*RAB27A*), and active regulation (*TNFRSF9, LAG3*), confirming that dominant TCR-expressing CD8 Tconv are the clonally-expanded antigen-driven cytotoxic cells in the skin, which are enriched in blister fluid. Intriguingly, some CD8+ Tconv cells expressed more than one TCRα/β chain. For example, the two dominantly-expanded TCRs with the same CASS<u>P</u>DRGGYEQYF CDR3β were predominantly expressed with a second CDR3α on the same CD8+ Tconv cells in blister fluid (222 cells, Extended Data Fig. 5i-ii). Cells expressing the third clonotype did not express a second TCR. These data were mirrored in affected skin (Extended Data Fig. 5ii-iii); suggesting a role for dual TCRαβ+ T-cells^14^ or two or more PIC (T-cells)^12^ in the pathogenesis of SJS/TEN.

Finally, we assessed the total clonality of ‘cytotoxic’ CD8+ Tconv cluster 3, where the three dominantly-expanded TCR showed similar representation among the top 50 clonotypes in affected skin (48%) and blister fluid (average 51%, Fig. 2Civ). In the affected skin, while up to 42% of cells expressed expanded clonotypes, >50% expressed unexpanded TCR (n=1 count, Fig. 2Cv). To investigate whether T-cells with unexpanded clonotypes had a distinct phenotype, we used blister fluid to increase the power for DEG analyses. Expanded oligoclonal clonotypes were significantly enriched for cytotoxicity-related genes, including *GNLY, PRF1*, and *FCGR3A*, but unexpanded polyclonal clonotypes for *GZMA* and *GZMK* (Fig. 2Di). Compared to TCR+ cells from inactivated cluster 2, unexpanded polyclonal clonotypes in cluster 3 remained significantly enriched for *GNLY, PRF1*, and *FCGR3A*, albeit less than expanded oligoclonal clonotypes, which were uniquely enriched for markers of activation (*TNFRSF9*), long intergenic non-coding (linc)RNAs (*LINC01871, LINC01943*), and immunomodulatory genes (*TIGIT, KLRC1, KLRD1*, Fig. 2Dii).

In CD8+ Tconv cluster 3 of SJS/TEN blister fluids, while unexpanded T-cells expressed diverse CDR3αβ, many expressed the same TRBV27/TRBJ2-7 of dominantly-expanded clonotypes (Extended data Fig. 6). In fact, several polyclonal unexpanded T-cells expressed the dominantly-expanded CDR3β (CASS<u>P</u>DRGGYEQYF) or had one amino acid mismatch (P to H/V/Y/F/L). Again, these clonotypes were expressed on single-cells expressing two TCRs (Extended Data Table 4), and GLIPH2 predicted shared peptide binding (S_DRGGYE CDR3β motif, fisher p=7.90E-15). Notably, the dominantly-expanded CASS<u>Q</u>DRGGYEQYF CDR3β was not identified on polyclonal T-cells, and the polyclonal T-cells with a shared CDR3β were only observed in blister fluids and not affected skin, suggesting incomplete capture of the TCR repertoire. Indeed, between blister fluids, only up to 10% of polyclonal T-cells in CD8 Tconv cluster 3 shared a TCR, raising the possibility that at least some had not been triggered by antigen-specific TCRs but site-specific ‘bystander’ activation^15^. However, both oligoclonal and polyclonal T-cells significantly downregulated *IL7R* compared to those in control cluster 2, indicative of TCR-dependent activation, and remained *KLRK1*^LO^, for which upregulation is associated with TCR-independent activation^15^ (Fig. 2Diii). The downregulation of IL7R was mirrored by scCITE-seq expression (CD127, data not shown).

Using a multiomic approach, we define the composition of single-cells at the site of SJS/TEN tissue damage that demonstrate blister fluid as an immune-rich reservoir of clonally-relevant cytotoxic CD8+ T-cells. Limitations are that, unlike the blister fluid, the composition of skin is likely to be affected by enzymatic digestion, and despite qualitative capture of expanded clonotypes, the power to detect unexpanded clonotypes is incomplete. We could also not control all patient-specific factors such as severity and time since onset of symptoms. However, we characterize the same cytotoxic CD8+ subpopulation, which expressed the same expanded TCRαβ clonotypes in affected skin and blister fluids but not in unaffected skin. Expanded clonotypes were enriched for *KLRD1, KLRC1, TNFRSF9*, and *FCGR3A*, indicative of terminally-differentiated effectors driving antigen-specific response^16-18^. Indeed, although *KLRC1* and *KLRD1* form the inhibitory NKG2A/CD94 receptor, their expression on otherwise activated T-cells is shown only to limit excessive activation to sustain immune response^18^. Further, expanded clonotypes are expressed (i) under influence of *LINC01871*, associated with TCR-dependent cytotoxicity in Sjogren’s disease^19^, (ii) on a subset of T-cells expressing two TCRαβ, associated with auto-and allo-cross-reactivities^14^, and (iii) in the same pathogenic cluster as diverse polyclonal T-cells, with a more limited but nonetheless cytolytic signature consistent with antigen-induced activation^15^. These data support a ‘selective-signaling’ model of TCR-triggered activation whereby different clonotypes support proliferation and/or cytolytic secretion. These functional differences can be observed by introducing molecular changes in antigens to stimulate the same TCR^20^, indicating that different TCR-ligand affinities determine how the T-cell will respond^21^.

Our data support the hypothesis that while drug-expanded oligoclonal CD8+ T-cells are highly cytotoxic, antigen-reactive polyclonal CD8+ T-cells also contribute to the effector pathogenesis, and the total number and affinity of cytotoxic TCR may impact disease severity. However, while oligoclonal CD8+ T-cells have been identified as drug-reactive^3,13,22^, polyclonal CD8+ T-cells may be driven by distinct drug-or self-antigen. Thus, as we define private oligoclonal and polyclonal TCRs in cytotoxic Tconv cluster 3 across patients with diverse drug-induced SJS/TEN, this dataset provides a multiomic atlas of antigen-driven T-cells clustered by drug and population that will help define cell-cell and receptor-ligand interactions towards tissue-relevant biological markers for earlier diagnosis and treatment. Further, the identification of all cytotoxic TCR clonotypes will inform functional assays to define complete HLA-and epitope-specificities towards novel genetic and structural prevention strategies.

## Methods

### Sample acquisition and storage

Burn blister fluid and healthy skin sections from the discarded edges of cutaneous surgeries were obtained from the Western Australian state burns unit. Acute SJS/TEN skin biopsies and aspirated blister fluid were obtained from cryopreserved biorepositories of clinically-confirmed SJS/TEN patients in the US (Vanderbilt University Medical Centre), South Africa (University of Cape Town), and Australia (Austin Health and University of Melbourne). SJS/TEN patients varied in age (range, 18-74 years; median 40 years), sex (male, n=6; female n=11), underlying disease, culprit drug, reaction timeline, genetic race, and clinical treatment. Briefly, upon immediate collection after excision, skin biopsies were cut into sub-centimeter pieces and placed into cryogenic tubes containing 80% fetal bovine serum (FBS) and 20% DMSO and flash-frozen at −80°C before being transferred into liquid nitrogen for long-term storage. Blister fluid samples were strained through a 70μm mesh filter and washed twice with PBS. The resulting cell pellet was resuspended in 90% FBS 10% DMSO freeze media and placed into a slow-cooling container (Mr. Frosty) at −80°C before being transferred into liquid nitrogen for long-term storage. All samples, clinical data, and analyses were obtained with informed consent and state health and local institutional review board (IRB) approval (Western Australian Department of Health RGS0000001924, Murdoch HREC 2011/056, 2017/246, and 2019/153). No compensation was provided to any participant for donated tissue or blister fluid, and all samples and data were de-identified.

### Enzymatic digestion of skin into a single-cell suspension

Tissue including 4mm punch biopsies from SJS/TEN patients and healthy skin from unrelated donors was quickly thawed in a water bath (37°C) and subjected to enzymatic digestion to obtain a single-cell suspension using published methods optimized for lymphoid cell recovery^23^ and single-cell sequencing^24^. Briefly, each sample was washed twice in PBS immediately after thawing to remove freezing media and resuspended in RPMI 1640 medium with 10% fetal bovine serum (FBS), 1mg/ml collagenase P and 40ug/ml DNAase I with agitation for 90 minutes. Tissue lysates were washed twice in PBS and passed through a 70μm Flowmi™ Cell Strainer. Viable cells were counted using trypan blue and an automated cell counter (Countess, Invitrogen). Cells were immediately resuspended in 22.5μl cell staining buffer for single-cell analyses.

### Surface antibody staining protocol for scCITE-sequencing

Single-cell suspensions of thawed and enzymatically-digested skin or unprocessed blister fluid were subject to surface antibody staining for a panel of up to 137 functional and surface lineage markers in the TotalSeq™-C Human Universal Cocktail, V1.0 (Biolegend) as per manufacturer’s protocol

(https://www.biolegend.com/Files/Images/media_assets/support_protocol/20-0014-00_TotalSeq-UC_8x11.pdf). Briefly, lyophilized antibody panels were resuspended in cell staining buffer (Biolegend Cat. No. 420201), vortexed, and incubated for 5 mins at room temperature. Antibodies were again vortexed and centrifuged (14,000xg, 10 mins, 4°C) while cell suspensions were blocked using 2.5μl Human TruStain FcX™ Fc blocking reagent and incubated for 10mins at 4°C. 25μl of reconstituted antibody cocktail was then added to 25μl of FcR-blocked cells, and for samples to be pooled into the same well of the 10x chromium chip, 1μl of the appropriate hashtag antibody was also added, with subsequent incubation for 30mins at 4°C. Each cell-antibody cocktail was then resuspended in 3.5ml cell staining buffer and washed three times in PBS at 400 x g for 5 mins at 4°C before cells were passed through a 70μm Flowmi™ Cell Strainer. Cell count and viability were recorded using trypan blue and an automated cell counter and cells were diluted as per the manufacturer’s instruction for loading onto the 10X Chromium Next GEM chip K.

### Single-cell 5’ RNA-TCR-CITE-sequencing

Cells were loaded onto the chromium Next GEM chip K chip at the manufacturer (10x Genomics) recommended concentration and single-cell libraries made using a chromium controller and Chromium Next GEM Single Cell 5’ Reagent Kit, Version 2.0 (10x Genomics) as per manufacturer instruction (https://www.10xgenomics.com/support/single-cell-immune-profiling/documentation/steps/library-prep/chromium-single-cell-5-reagent-kits-user-guide-v-2-chemistry-dual-index-with-feature-barcoding-technology-for-cell-surface-protein-and-immune-receptor-mapping), with a target of 1,000-20,000 cells. 5’ libraries were then sequenced using the Illumina NovaSeq 6000 platform for the capture of 70,000 reads/cell, including 50,000 for cDNA, 5,000 for TCR, and 15,000 for the cell surface protein library. Sample sequencing was completed in the VANderbilt Technologies for Advanced GEnomics core (VANTAGE), with subsequent bioinformatic and computational analyses at IIID, Murdoch University.

### Computational and statistical analysis

The single-cell sequencing data were processed using CellRanger v6.1.2 (10x Genomics) to demultiplex individual cell barcodes, remove PCR duplicates and align RNA reads to the reference GRCh38 human transcriptome. The individual hashes within each 10x run were further demultiplexed using Souporcell^25^ to cluster samples based on SNPs in the RNA-seq reads. The hashtag negatives or doublets, as well as cells with low (<500) unique molecular identifiers (UMIs), were removed. Cells with <100 genes and >50% mitochondrial content were removed to filter low-quality, dead, or dying populations, with a second pass filter excluding apoptotic populations characterized by a low percentage of ribosomal genes versus a high percentage of mitochondrial genes, using a defined mitochondrial-ribosomal RNA ratio of >0.47^26^. Downstream transcriptome-based graph clustering and principal component analyses (PCA) with cell phenotype consensus calling were performed using R Seurat v4.1.1 package^9^, without input of TCR, BCR, mitochondrial, ribosomal, or sex-linked genes to avoid bias due to known expression variability. Subsets with <50 scRNA-defined cells were removed. While most cell subsets formed distinct clusters, keratinocytes predominantly aligned to a shared cluster including a minority of T-cells, NK cells, and monocytes at the UMAP origin defined by lower UMI; indicative of cell stress. However, these cells met all other criteria for inclusion and were retained for pathogenic importance in the context of drug-induced stress during SJS/TEN. Batch correction for 10x run, sample type (skin or blister fluid), and cell cycle phase was performed using harmony^27^, and experimental doublets were identified and removed using a majority consensus benchmark^28^ of three independent bioinformatic algorithms; DoubletFinder^29^, scDBLfinder^30^, scDS^31^. T and NK subsets were re-aligned to the HuBMAP consortium Azimuth PBMC reference multimodal dataset^9^ for consensus calling of conventional and unconventional subsets. Visualization and differential expression analyses were performed using Visual Genomics Analysis Studio (VGAS)^32^. Differential gene expression analyses were performed using the Wilcoxon rank-sum test with Benjamini-Hochberg adjusted correction (adj. p<0.05). Pathway analyses were performed using Enrichr^33^. To avoid the bias of testing compositional changes of each cell type independently using univariate models, statistical analysis of single-cell composition was performed using scCODA^34^ and the software-recommended standard and minimum significance thresholds of p<0.05 and p<0.4, respectively.

## Acknowledgments

We gratefully acknowledge patients and their families, the significant expertise of the VANderbilt Technologies for Advanced GEnomics core (VANTAGE) core for sample sequencing, and researchers and clinicians from the Australasian Registry for Severe Cutaneous Adverse Reaction (AUS-SCAR) and African Registry of Severe Cutaneous Adverse Reactions (AFRI-SCAR) for the provision of clinically-curated SJS/TEN tissue samples from Australasia and South Africa, respectively. These thanks are extended but not limited to (AUS-SCAR) Jason A Trubiano, Fiona James, Ar Kar Aung, Michelle S Y Goh, Celia Zubrinich, Douglas Gin, Heather Cleland, Andrew Awad, (AFRI-SCAR) Jonny G Peter, Rannakoe J Lehloenya, Phuti Choshi, Sarah Pedretti, Tafadzwa Chimbetete, Rose Selim, and Mireille Porter. Further, for the provision of control samples, we are grateful to and acknowledge Mark Fear and Fiona M Wood (Burn Injury Research Unit, School of Surgery, University of Western Australia, Australia; The Burns Service of WA, WA Department of Health, Murdoch, Australia), and Allison Hanlon (Dept of Medicine, Vanderbilt University Medical Centre, Nashville, TN, USA).

## Data Availability

All data produced in the present study will be available after peer-review, when published.

