## Extended Data Figure 1 for "Multiomic single-cell sequencing defines tissue-specific responses in Stevens-Johnson Syndrome and Toxic epidermal necrolysis"

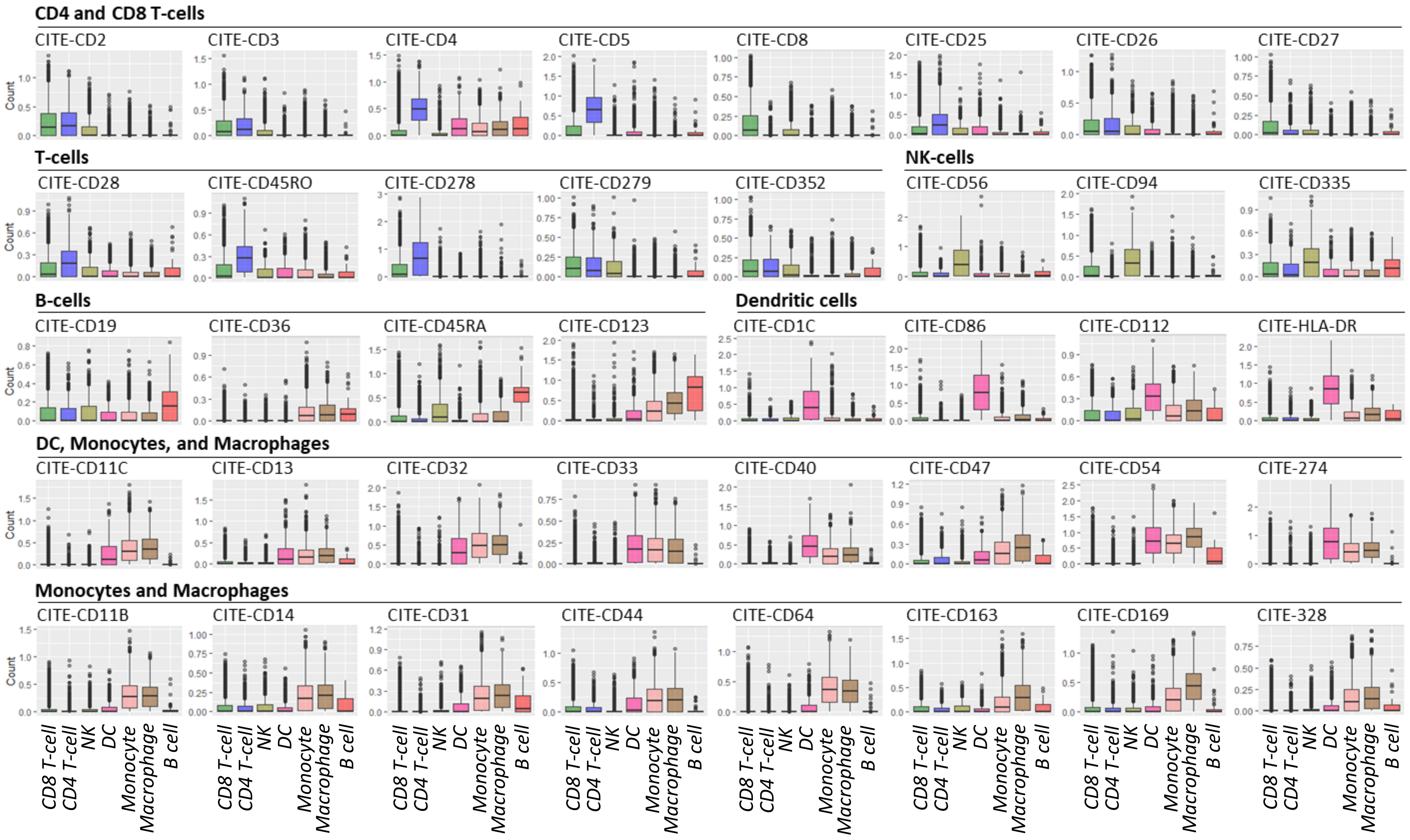


**Extended data Figure 1. Cell surface protein expression (scCITE-seq) of key lineage and activation markers on scRNA-defined immune cell subsets.** Box plots show the median expression for each cell surface protein across scRNA-subsets from a representative UMAP of SJS/TEN blister fluid and 17,629 cells obtained within a single 10x run. Surface proteins are grouped into panels headed by the scRNA-defined cell type with the highest median expression for that marker.
