## Extended Data Figure 2 for "Multiomic single-cell sequencing defines tissue-specific responses in Stevens-Johnson Syndrome and Toxic epidermal necrolysis"

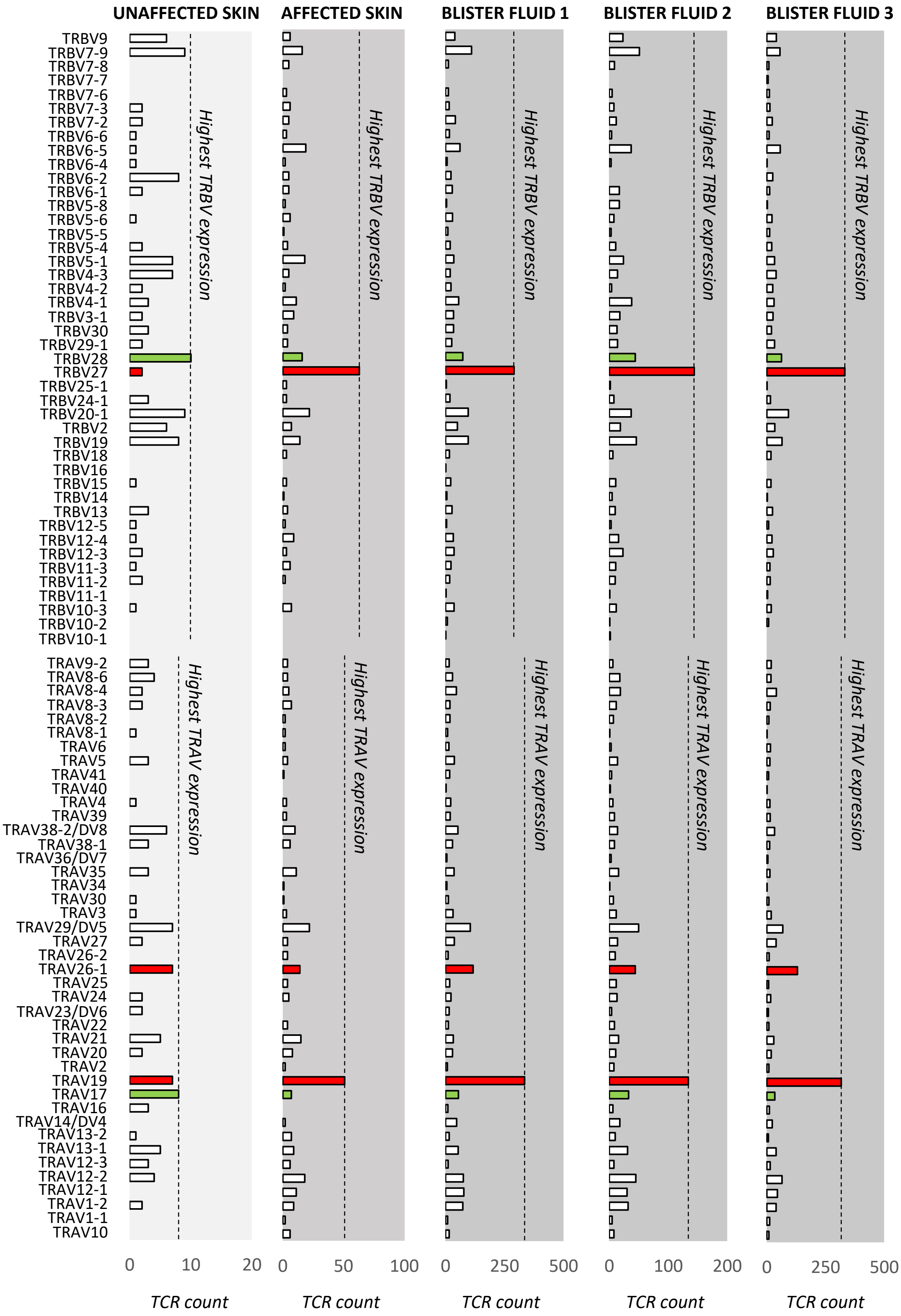


**Extended data Figure 2. Individual TRAV and TRBV scTCR-seq counts on CD8 Tconv cells across paired samples from a single SJS/TEN patient.** Individual counts for all identified TRAV and TRBV in CD8 Tconv across samples from unaffected skin, affected skin, and blister from three anatomical sites. The dominant TRAV and TRBV in affected skin and blister fluid are highlighted red, and the dominant TRAV and TRBV in unaffected skin are highlighted green. Figure created using VGAS. *TCR, T-cell receptor; TRAV, TCR alpha variable; TRBV, TCR beta variable; Tconv, T conventional cell.*
