## Extended Data Figure 3 for "Multiomic single-cell sequencing defines tissue-specific responses in Stevens-Johnson Syndrome and Toxic epidermal necrolysis"

**(i)** **UNAFFECTED SKIN (79 CD8+ Tconv cells with TCR expression)**

| TCR | CDR3α | CDR3β | TRAV | TRAJ | TRBV | TRBJ | Ct. |
|---|---|---|---|---|---|---|---|
| 1 | CIVRVARNTGNQFYF | CASSYHTGGSGYEQYF | TRAV26-1 | TRAJ49 | TRBV6-2 | TRBJ2-7 | 3 |
| 2 | CATDVMNRDDKIIF | CASSLTARGSGVELFF | TRAV17 | TRAJ30 | TRBV13 | TRBJ2-2 | 2 |
| 3 | CAVSDRSGGGADGLTF | CASSLGLNYEQYF | TRAV8-6 | TRAJ45 | TRBV28 | TRBJ2-7 | 2 |
| 4 | CAASVTGNQFYF | CASSVESGGWYGYTF | TRAV29/DV5 | TRAJ49 | TRBV9 | TRBJ1-2 | 2 |
| 5 | CALRGWGRRALTF | CATSDDGAGTDTQYF | TRAV19 | TRAJ5 | TRBV24-1 | TRBJ2-3 | 1 |
| 6 | CAGAGNAGNMLTF | CASSQVRFYGYTF | TRAV27 | TRAJ39 | TRBV4-3 | TRBJ1-2 | 1 |
| 7 | CALTQGAQKLVF | CASSHLTGELFF | TRAV16 | TRAJ54 | TRBV11-2 | TRBJ2-2 | 1 |
| 8 | CAETSYGQNFVF | CASSEMSVGETQYF | TRAV5 | TRAJ26 | TRBV6-1 | TRBJ2-5 | 1 |
| 9 | CALRGGAAGNKLTF | CASSDNPPYTEAFF | TRAV19 | TRAJ17 | TRBV7-9 | TRBJ1-1 | 1 |
| 10 | CALSEVTTSGTYKYIF | CASSQDRGGYEQYF | TRAV19 | TRAJ40 | TRBV27 | TRBJ2-7 | 1 |

**AFFECTED SKIN (235 CD8+ Tconv cells with TCR expression)**

| TCR | CDR3α | CDR3β | TRAV | TRAJ | TRBV | TRBJ | Ct. |
|---|---|---|---|---|---|---|---|
| 1 | CALSEVTTSGTYKYIF | CASSQDRGGYEQYF | TRAV19 | TRAJ40 | TRBV27 | TRBJ2-7 | 27 |
| 2 | CALSEVTTSGTYKYIF | CASSPDRGGYEQYF | TRAV19 | TRAJ40 | TRBV27 | TRBJ2-7 | 20 |
| 3 | CIVRVHSGGGADGLTF | CASSPDRGGYEQYF | TRAV26-1 | TRAJ45 | TRBV27 | TRBJ2-7 | 10 |
| 4 | CVVNNARNNDMRF | CASSPDRGGYEQYF | TRAV12-1 | TRAJ43 | TRBV27 | TRBJ2-7 | 4 |
| 5 | CLNDMRF | CASSQLSGNSPLHF | TRAV25 | TRAJ43 | TRBV3-1 | TRBJ1-6 | 3 |
| 6 | CAGRPDSGTYKYIF | CPPSLPRDDYEQYF | TRAV35 | TRAJ40 | TRBV27 | TRBJ2-7 | 3 |
| 7 | CAVCQEDDYKLSF | CSARDLAVYNSPLHF | TRAV22 | TRAJ20 | TRBV20-1 | TRBJ1-6 | 2 |
| 8 | CAASVTGNQFYF | CASSVESGGWYGYTF | TRAV29/DV5 | TRAJ49 | TRBV9 | TRBJ1-2 | 2 |
| 9 | CAVSGYGGATNKLIF | CASSLGDRQSYEQYF | TRAV21 | TRAJ32 | TRBV7-9 | TRBJ2-7 | 2 |
| 10 | CAVSPNNNARLMF | CASSLLGVGSPLHF | TRAV21 | TRAJ31 | TRBV5-1 | TRBJ1-6 | 2 |

**BLISTER FLUID 1 (ARM, 1336 CD8+ Tconv cells with TCR expression)**

| TCR | CDR3α | CDR3β | TRAV | TRAJ | TRBV | TRBJ | Ct. |
|---|---|---|---|---|---|---|---|
| 1 | CALSEVTTSGTYKYIF | CASSPDRGGYEQYF | TRAV19 | TRAJ40 | TRBV27 | TRBJ2-7 | 129 |
| 2 | CIVRVHSGGGADGLTF | CASSPDRGGYEQYF | TRAV26-1 | TRAJ45 | TRBV27 | TRBJ2-7 | 98 |
| 3 | CALSEVTTSGTYKYIF | CASSQDRGGYEQYF | TRAV19 | TRAJ40 | TRBV27 | TRBJ2-7 | 82 |
| 4 | CALSEARSSASKIIF | CASSDRDRYYEQYF | TRAV19 | TRAJ3 | TRBV7-9 | TRBJ2-7 | 33 |
| 5 | CVVNNARNNDMRF | CASSPDRGGYEQYF | TRAV12-1 | TRAJ43 | TRBV27 | TRBJ2-7 | 24 |
| 6 | CALSESETSGSRLTF | CASSLWEVERAYNEQFF | TRAV19 | TRAJ58 | TRBV28 | TRBJ2-1 | 18 |
| 7 | CAVSLTYSGGGADGLTF | CSAKGGEQYF | TRAV8-4 | TRAJ45 | TRBV20-1 | TRBJ2-7 | 17 |
| 8 | CAADTGGFKTIF | CASTLSAGLNQPQHF | TRAV13-1 | TRAJ9 | TRBV19 | TRBJ1-5 | 16 |
| 9 | CVVNLYKLSF | CASSSQRAVDEQFF | TRAV12-1 | TRAJ20 | TRBV7-9 | TRBJ2-1 | 14 |
| 10 | CATGTSYGKLTF | CASSLPTLGLAGGATDNEQFF | TRAV17 | TRAJ52 | TRBV28 | TRBJ2-1 | 13 |

**BLISTER FLUID 2 (FACE, 616 CD8+ Tconv cells with TCR expression)**

| TCR | CDR3α | CDR3β | TRAV | TRAJ | TRBV | TRBJ | Ct. |
|---|---|---|---|---|---|---|---|
| 1 | CALSEVTTSGTYKYIF | CASSQDRGGYEQYF | TRAV19 | TRAJ40 | TRBV27 | TRBJ2-7 | 43 |
| 2 | CALSEVTTSGTYKYIF | CASSPDRGGYEQYF | TRAV19 | TRAJ40 | TRBV27 | TRBJ2-7 | 42 |
| 3 | CIVRVHSGGGADGLTF | CASSPDRGGYEQYF | TRAV26-1 | TRAJ45 | TRBV27 | TRBJ2-7 | 29 |
| 4 | CALSEARSSASKIIF | CASSDRDRYYEQYF | TRAV19 | TRAJ3 | TRBV7-9 | TRBJ2-7 | 14 |
| 5 | CAMNSYSGAGSYQLTF | CASSPFYSGGDTDTQYF | TRAV14/DV4 | TRAJ28 | TRBV12-3 | TRBJ2-3 | 9 |
| 6 | CAADTGGFKTIF | CASTLSAGLNQPQHF | TRAV13-1 | TRAJ9 | TRBV19 | TRBJ1-5 | 9 |
| 7 | CATGTSYGKLTF | CASSLPTLGLAGGATDNEQFF | TRAV17 | TRAJ52 | TRBV28 | TRBJ2-1 | 7 |
| 8 | CAVSLTYSGGGADGLTF | CSAKGGEQYF | TRAV8-4 | TRAJ45 | TRBV20-1 | TRBJ2-7 | 6 |
| 9 | CALSESETSGSRLTF | CASSLWEVERAYNEQFF | TRAV19 | TRAJ58 | TRBV28 | TRBJ2-1 | 6 |
| 10 | CVVNLYKLSF | CASSSQRAVDEQFF | TRAV12-1 | TRAJ20 | TRBV7-9 | TRBJ2-1 | 6 |

**BLISTER FLUID 3 (FOOT, 1039 CD8+ Tconv cells with TCR expression)**

| TCR | CDR3α | CDR3β | TRAV | TRAJ | TRBV | TRBJ | Ct. |
|---|---|---|---|---|---|---|---|
| 1 | CALSEVTTSGTYKYIF | CASSPDRGGYEQYF | TRAV19 | TRAJ40 | TRBV27 | TRBJ2-7 | 154 |
| 2 | CIVRVHSGGGADGLTF | CASSPDRGGYEQYF | TRAV26-1 | TRAJ45 | TRBV27 | TRBJ2-7 | 112 |
| 3 | CALSEVTTSGTYKYIF | CASSQDRGGYEQYF | TRAV19 | TRAJ40 | TRBV27 | TRBJ2-7 | 93 |
| 4 | CAVSLTYSGGGADGLTF | CSAKGGEQYF | TRAV8-4 | TRAJ45 | TRBV20-1 | TRBJ2-7 | 18 |
| 5 | CAVYYGNNRLAF | CASSTGGLGNQPQHF | TRAV12-2 | TRAJ7 | TRBV6-5 | TRBJ1-5 | 14 |
| 6 | CVVNNARNNDMRF | CASSPDRGGYEQYF | TRAV12-1 | TRAJ43 | TRBV27 | TRBJ2-7 | 11 |
| 7 | CATGTSYGKLTF | CASSLPTLGLAGGATDNEQFF | TRAV17 | TRAJ52 | TRBV28 | TRBJ2-1 | 9 |
| 8 | CAMNSYSGAGSYQLTF | CASSPFYSGGDTDTQYF | TRAV14/DV4 | TRAJ28 | TRBV12-3 | TRBJ2-3 | 9 |
| 9 | CAATGSGTYKYIF | CASSMQGYTMNTEAFF | TRAV29/DV5 | TRAJ40 | TRBV19 | TRBJ1-1 | 8 |
| 10 | CALSEARSSASKIIF | CASSDRDRYYEQYF | TRAV19 | TRAJ3 | TRBV7-9 | TRBJ2-7 | 7 |

**(ii)** **UNAFFECTED SKIN** **AFFECTED SKIN** **BLISTER FLUID 1** **BLISTER FLUID 2** **BLISTER FLUID 3**

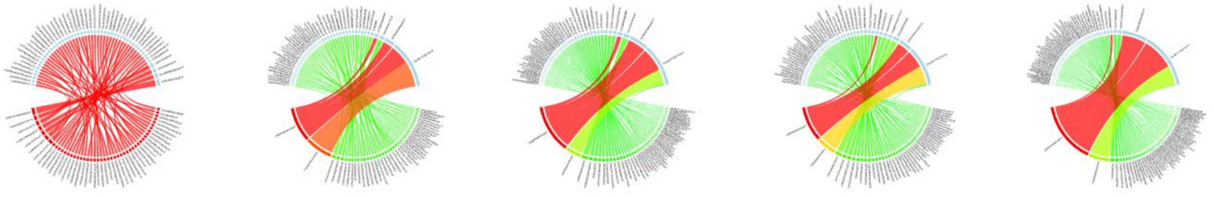

**Extended Data Figure 3. Top functional CDR3αβ clonotypes and counts in CD8 Tconv across paired samples from a single SJS/TEN patient.** (i) Top 10 CDR3 TCRαβ clonotypes and counts in the CD8 Tconv population. Blue highlights indicate the same functional CDR3αβ clonotypes. (ii) Circos plots show comparative clonality and CDR3α and CDR3β pairings between samples. The width of each segment is proportionate to its expression. Least to most dominant is colored green to red. Top 50 CDR3αβ clonotypes shown for each sample. *Tconv, T conventional cell; TCR, T-cell receptor; TRAV, TCR alpha variable; TRBV, TCR beta variable; TRAJ, TCR alpha joining; TRBJ, TCR beta joining; CDR3, complementary-determining region; Ct., count.*
