## Extended Data Figure 4 for "Multiomic single-cell sequencing defines tissue-specific responses in Stevens-Johnson Syndrome and Toxic epidermal necrolysis"

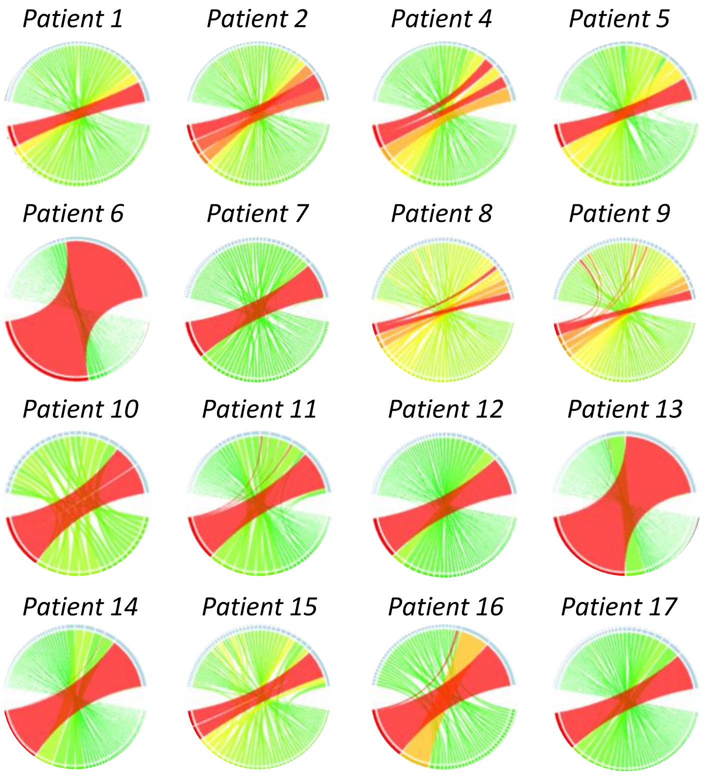


**Extended Data Figure 4. Oligoclonal TCR CDR3αβ clonotypes were identified in CD8+ Tconv populations in blister fluids from all SJS/TEN patients.** Circos plots show comparative clonality and CDR3α and CDR3β pairings between samples. The width of each segment is proportionate to its expression. Least to most dominant is colored green to red. Up to the top 50 clonotypes are shown for each sample. Patient 3 is excluded from the panel as already depicted in Figure 2. Tconv, T conventional cell; TCR, T-cell receptor; CDR, complimentary-determining region.
