## Extended Data Figure 5 for "Multiomic single-cell sequencing defines tissue-specific responses in Stevens-Johnson Syndrome and Toxic epidermal necrolysis"

**(i) Blister fluid: Expression of top expanded TCRs in CD8 Tconv**

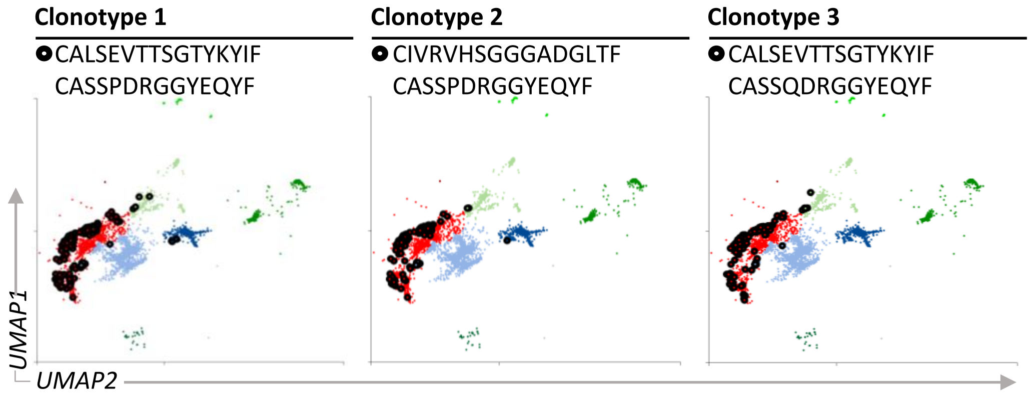


**(ii) Blister fluid: CDR3αβ expression of CD8 Tconv cells selected by each top clonotype**

| | **Cells expressing clonotype 1** | | **Cells expressing clonotype 2** | | **Cells expressing clonotype 3** | |
|---|---|---|---|---|---|---|
| **TCR** | **CDR3α/CDR3β** | **Ct.** | **CDR3α/CDR3β** | **Ct.** | **CDR3α/CDR3β** | **Ct.** |
| 1 | CALSEVTTSGTYKYIF<br>CASSPDRGGYEQYF | 325 | CIVRVHSGGGADGLTF<br>CASSPDRGGYEQYF | 239 | CALSEVTTSGTYKYIF<br>CASSQDRGGYEQYF | 218 |
| 2 | CIVRVHSGGGADGLTF<br>CASSPDRGGYEQYF | 222 | CALSEVTTSGTYKYIF<br>CASSPDRGGYEQYF | 222 | | |
| 3 | CVVNNARNNDMRF<br>CASSPDRGGYEQYF | 37 | CIVRVHSGGGADGLTF<br>CASTLSAGLNQPQHF | 1 | | |
| 4 | CALSEVTTSGTYKYIF<br>CASSYDRGGYEQYF | 1 | CAADTGGFKTIF<br>CASSPDRGGYEQYF | 1 | | |
| 5 | | | CAADTGGFKTIF<br>CASTLSAGLNQPQHF | 1 | | |

**(iii) Affected skin: CDR3αβ expression of CD8 Tconv cells selected by each top clonotype**

| | **Cells expressing clonotype 1** | | **Cells expressing clonotype 2** | | **Cells expressing clonotype 3** | |
|---|---|---|---|---|---|---|
| TCR | CDR3α/CDR3β | Ct. | CDR3α/CDR3β | Ct. | CDR3α/CDR3β | Ct. |
| 1 | CALSEVTTSGTYKYIF<br>CASSPDRGGYEQYF | 20 | CALSEVTTSGTYKYIF<br>CASSPDRGGYEQYF | 10 | CALSEVTTSGTYKYIF<br>CASSQDRGGYEQYF | 27 |
| 2 | CIVRVHSGGGADGLTF<br>CASSPDRGGYEQYF | 10 | CIVRVHSGGGADGLTF<br>CASSPDRGGYEQYF | 10 | | |
| 3 | CVVNNARNNDMRF<br>CASSPDRGGYEQYF | 4 | | | | |

**Extended Data Figure 5. Top expanded TCRαβ clonotypes in CD8 Tconv in affected skin and blister fluid have shared expression on dual TCRαβ+ T-cells.** (i) Expression of each of the top 3 dominantly-expanded TCR on CD8 Tconv cells in blister fluid from a patient with SJS/TEN. Cells expressing each clonotype are circled in black. (ii) The TCR clonotypes and counts expressed by cells selected for expression of each top clonotype individually in SJS/TEN blister fluid. (iii) The TCR clonotypes and counts expressed by cells selected for expression of each top clonotype individually in SJS/TEN affected skin. Grey highlight indicates TCR expressed by the same cells. Ct., cell count. *Tconv, T conventional cell; TCR, T-cell receptor; CDR, complimentary-determining region; Ct., Count.*
