## Extended Data Figure 6 for "Multiomic single-cell sequencing defines tissue-specific responses in Stevens-Johnson Syndrome and Toxic epidermal necrolysis"

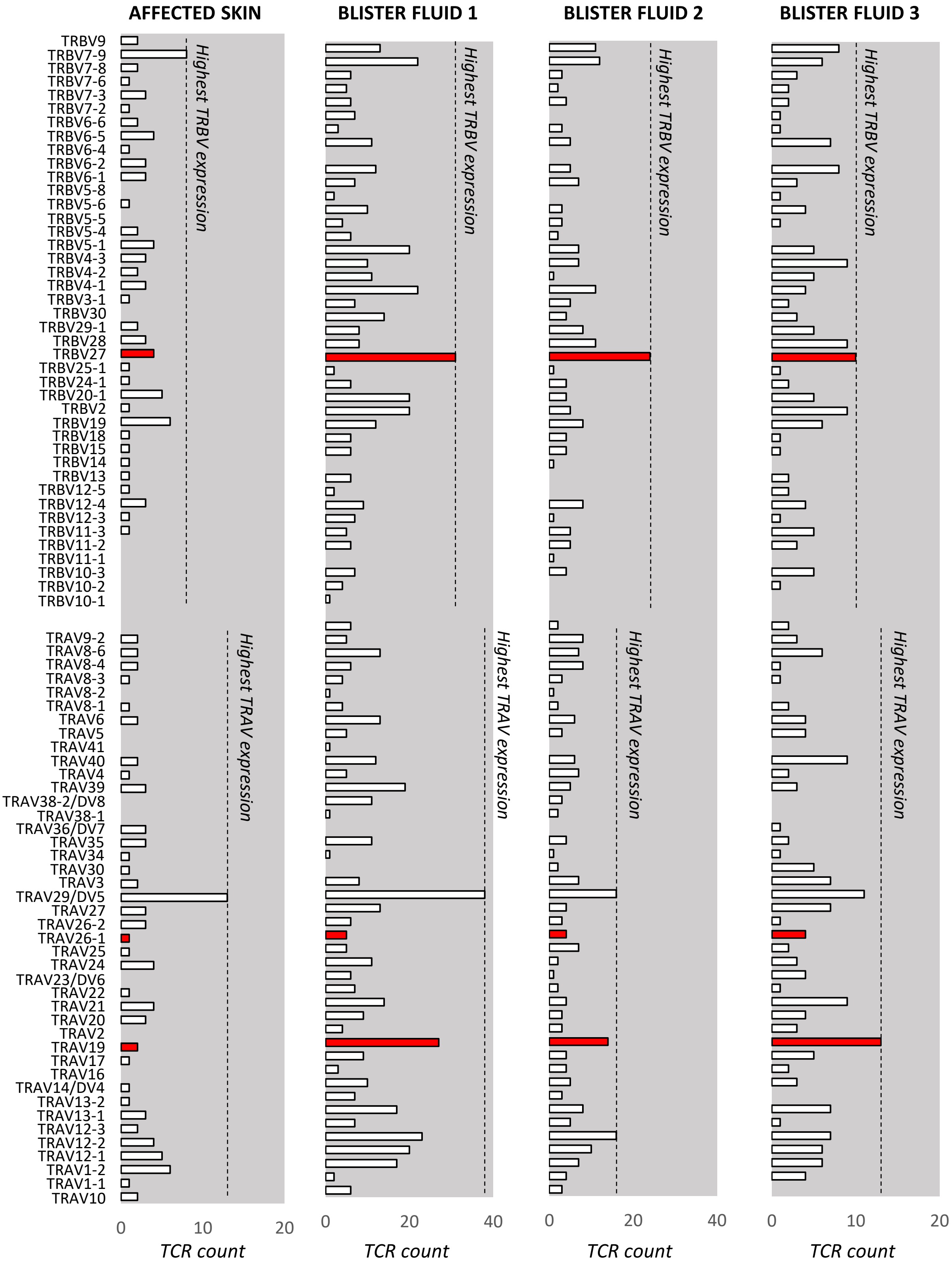


**Extended data Figure 6. Individual TRAV and TRBV scTCR-seq counts on unexpanded cells of CD8+ Tconv cluster 3 across samples from a single SJS/TEN patient.** Individual counts for all identified TRAV and TRBV genes across samples from unaffected skin, affected skin, and blister from three anatomical sites. The dominant TRAV and TRBV of the dominantly-expanded cells in CD8+ Tconv cluster 3 in affected skin and blister fluid are highlighted in red. Figure created using VGAS. *TCR, T-cell receptor; TRAV, TCR alpha variable; TRBV, TCR beta variable; Tconv, T conventional cell.*
