## Extended Data Table 1 for "Multiomic single-cell sequencing defines tissue-specific responses in Stevens-Johnson Syndrome and Toxic epidermal necrolysis"

**Extended Data Table 1. Differential gene signatures of Seurat-defined CD8 Tconv clusters 1-8.** The top 25 genes that show significant (Wilcoxon rank sum test, Benjamini Hochberg adjusted p <0.05) increased expression are shown in order of fold change. Red font indicates genes below significance. Genes shared or indicative of shared functional state between clusters 1-2 (blue), clusters 3-4 (red), and clusters 5-8 (green) are highlighted. *Tconv, T conventional cell; FC, fold change.*

| | Quiescent/inactive | | | | | | Cytotoxic/proliferative | | | | | |
|---|---|---|---|---|---|---|---|---|---|---|---|---|
| | CD8 Tconv cluster 1 | | | CD8 Tconv cluster 2 | | | CD8 Tconv cluster 3 | | | CD8 Tconv cluster 4 | | |
| Rank | Gene | Log2FC | p value | Gene | Log2FC | p value | Gene | Log2FC | p value | Gene | Log2FC | p value |
| 1 | DCD | 11.4 | 1.7E-04 | CMC1 | 2.7 | 2.2E-29 | TYMS | 4.8 | 4.0E-37 | HIST1H2AI | 4.9 | 7.7E-24 |
| 2 | MUCL1 | 6.7 | 1.5E-02 | LTB | 2.5 | 1.8E-25 | STMN1 | 3.3 | 5.4E-22 | MKI67 | 4.7 | 1.0E-22 |
| 3 | XIST | 1.9 | 6.2E-03 | FOS | 2.4 | 2.8E-23 | KLRC1 | 3.0 | 3.6E-19 | CENPF | 4.0 | 8.6E-15 |
| 4 | MALAT1 | 1.8 | 1.3E-02 | IL7R | 2.2 | 1.8E-19 | TRAV12-3 | 3.0 | 1.4E-18 | HIST1H4C | 3.3 | 5.0E-09 |
| 5 | NEAT1 | 1.7 | 2.0E-02 | DUSP1 | 1.9 | 7.4E-14 | GNLY | 2.9 | 2.2E-18 | HIST1H1E | 3.2 | 9.1E-09 |
| 6 | CTSD | 1.7 | 2.7E-02 | HSPA1A | 1.8 | 5.7E-11 | TRBV9 | 2.8 | 2.6E-17 | STMN1 | 3.1 | 6.5E-08 |
| 7 | GRK2 | 1.4 | 8.2E-02 | ZFP36 | 1.8 | 6.9E-13 | TUBA1B | 2.7 | 1.3E-16 | HIST1H1D | 3.0 | 2.8E-07 |
| 8 | MACF1 | 1.4 | 8.7E-02 | JUNB | 1.7 | 2.6E-11 | GALNT2 | 2.4 | 5.8E-14 | HIST1H1C | 2.8 | 3.8E-06 |
| 9 | AKNA | 1.4 | 9.0E-02 | DUSP2 | 1.6 | 5.1E-10 | TRBV4-1 | 2.1 | 3.0E-10 | CEBPD | 2.7 | 2.2E-05 |
| 10 | NLRC3 | 1.4 | 9.6E-02 | DNAJB1 | 1.5 | 2.1E-08 | TUBB | 2.0 | 4.6E-10 | LINC01871 | 2.2 | 3.7E-03 |
| 11 | SYNE2 | 1.4 | 9.5E-02 | TXNIP | 1.3 | 1.3E-06 | ACP5 | 2.0 | 5.5E-10 | GNLY | 2.2 | 4.2E-03 |
| 12 | NKTR | 1.4 | 1.0E-01 | CXCR4 | 1.3 | 1.5E-06 | HIST1H4C | 2.0 | 8.2E-10 | HMGB2 | 2.2 | 5.4E-03 |
| 13 | IKZF3 | 1.3 | 1.2E-01 | HLA-DPA1 | 1.3 | 2.6E-06 | HAVCR2 | 2.0 | 1.1E-09 | TUBA1B | 2.0 | 2.2E-02 |
| 14 | INPP5D | 1.3 | 1.2E-01 | HLA-DPB1 | 1.2 | 1.1E-05 | LGALS1 | 1.8 | 8.0E-09 | MT1E | 1.8 | 4.1E-02 |
| 15 | RNF213 | 1.3 | 1.2E-01 | ZFP36L2 | 1.1 | 3.2E-05 | GZMB | 1.8 | 1.3E-08 | IFI27 | 1.9 | 5.2E-02 |
| 16 | JUND | 1.3 | 1.3E-01 | CD6 | 1.1 | 3.5E-05 | HMGB2 | 1.8 | 1.6E-08 | ETS1 | 1.8 | 7.4E-02 |
| 17 | OGA | 1.3 | 1.3E-01 | GZMM | 1.1 | 3.8E-05 | FABP5 | 1.8 | 4.9E-08 | MT2A | 1.8 | 9.4E-02 |
| 18 | GNAS | 1.3 | 1.3E-01 | CD69 | 1.1 | 7.4E-05 | HMGN2 | 1.6 | 4.0E-07 | DUT | 1.7 | 9.9E-02 |
| 19 | MT-ND1 | 1.3 | 1.5E-01 | PDCD4 | 1.1 | 1.7E-04 | H2AFZ | 1.6 | 7.2E-07 | SET | 1.7 | 1.1E-01 |
| 20 | STK10 | 1.3 | 1.6E-01 | MALAT1 | 1.0 | 2.1E-04 | IFI27 | 1.5 | 1.9E-06 | H1FX | 1.7 | 1.4E-01 |
| 21 | MT-ND6 | 1.2 | 1.6E-01 | BTG1 | 1.0 | 2.8E-04 | MT1E | 1.5 | 2.0E-06 | H2AFV | 1.7 | 1.5E-01 |
| 22 | MT-ND2 | 1.2 | 1.5E-01 | JUN | 1.0 | 6.0E-04 | DUT | 1.5 | 2.5E-06 | ITGB1 | 1.7 | 1.5E-01 |
| 23 | KMT2A | 1.2 | 1.6E-01 | KLF6 | 1.0 | 7.0E-04 | PRF1 | 1.4 | 1.1E-05 | LYST | 1.5 | 2.1E-01 |
| 24 | MT-ND4 | 1.2 | 1.6E-01 | FTH1 | 0.9 | 2.9E-03 | PTMS | 1.4 | 1.2E-05 | MZT2B | 1.5 | 2.2E-01 |
| 25 | SRRM2 | 1.2 | 1.7E-01 | LINC02446 | 0.8 | 4.8E-03 | LAG3 | 1.4 | 2.3E-05 | IFI16 | 1.5 | 2.6E-01 |

| | Innate-like | | | | | | | | | | | |
|---|---|---|---|---|---|---|---|---|---|---|---|---|
| | CD8 Tconv cluster 5 | | | CD8 Tconv cluster 6 | | | CD8 Tconv cluster 7 | | | CD8 Tconv cluster 8 | | |
| Rank | Gene | Log2FC | p value | Gene | Log2FC | p value | Gene | Log2FC | p value | Gene | Log2FC | p value |
| 1 | S100A8 | 5.6 | 9.2E-47 | CST3 | 4.9 | 1.8E-11 | IDO1 | 7.5 | 6.0E-66 | TYROBP | 5.1 | 3.0E-15 |
| 2 | CST3 | 5.3 | 1.2E-41 | S100A8 | 4.5 | 4.6E-09 | CXCL10 | 7.4 | 1.2E-68 | XCL1 | 4.8 | 1.1E-08 |
| 3 | S100A9 | 5.1 | 2.2E-36 | TYROBP | 3.8 | 9.7E-06 | SERPING1 | 7.3 | 4.7E-66 | XCL2 | 4.6 | 4.3E-05 |
| 4 | TYROBP | 4.8 | 5.0E-31 | FCER1G | 3.3 | 6.4E-04 | IL1R2 | 7.3 | 9.4E-62 | FCER1G | 4.2 | 6.8E-08 |
| 5 | LYZ | 4.7 | 1.2E-27 | TRAV27 | 2.9 | 1.8E-02 | MARCKS | 7.2 | 4.3E-59 | KLRB1 | 4.0 | 7.3E-07 |
| 6 | FCER1G | 4.3 | 1.4E-22 | IFITM3 | 2.2 | 3.0E-01 | THBS1 | 7.0 | 3.2E-58 | CMC1 | 3.6 | 6.8E-05 |
| 7 | AIF1 | 3.6 | 3.7E-13 | FOS | 2.1 | 3.0E-01 | SERPINA1 | 6.9 | 3.8E-53 | AREG | 2.7 | 5.2E-02 |
| 8 | GLUL | 3.3 | 1.1E-10 | HIST1H2AI | 1.6 | 7.9E-01 | C15orf48 | 6.8 | 7.5E-52 | FOS | 2.5 | 9.8E-02 |
| 9 | FOS | 2.9 | 2.2E-07 | HLA-DPA1 | 1.3 | 1.0E+00 | SPI1 | 6.6 | 3.3E-48 | KLRD1 | 2.0 | 5.8E-01 |
| 10 | IFITM3 | 2.6 | 6.0E-06 | CTSB | 1.2 | 1.0E+00 | S100A8 | 6.6 | 1.1E-50 | DUSP1 | 2.0 | 4.3E-01 |
| 11 | HLA-DQA1 | 2.5 | 5.2E-05 | HIST1H1E | 1.2 | 1.0E+00 | CST3 | 6.6 | 5.2E-52 | PPP1R14B | 1.8 | 6.1E-01 |
| 12 | IFIT3 | 2.5 | 5.9E-05 | SAT1 | 1.2 | 1.0E+00 | CD14 | 6.5 | 4.0E-46 | IL7R | 1.8 | 9.1E-01 |
| 13 | HIST1H2AI | 2.4 | 1.7E-04 | FTL | 1.1 | 1.0E+00 | C1QA | 6.4 | 2.5E-46 | IFITM3 | 1.6 | 8.0E-01 |
| 14 | HIST1H1E | 2.0 | 3.4E-03 | HLA-DRA | 1.0 | 1.0E+00 | C1QB | 6.4 | 1.4E-45 | DUSP2 | 1.5 | 9.3E-01 |
| 15 | SAT1 | 1.9 | 9.9E-03 | TXN | 1.0 | 1.0E+00 | CTSL | 6.3 | 8.3E-40 | TXNIP | 1.5 | 1.0E+00 |
| 16 | HLA-DRA | 1.8 | 1.7E-02 | TNFSF10 | 0.9 | 1.0E+00 | C1QC | 6.3 | 5.7E-43 | ZFP36 | 1.2 | 1.0E+00 |
| 17 | JUND | 1.8 | 2.7E-02 | NEAT1 | 0.9 | 1.0E+00 | S100A9 | 6.1 | 4.8E-39 | BTG1 | 1.2 | 1.0E+00 |
| 18 | PARP14 | 1.7 | 4.5E-02 | TYMP | 0.9 | 1.0E+00 | TYROBP | 5.9 | 4.2E-37 | MBP | 1.1 | 1.0E+00 |
| 19 | HLA-DPA1 | 1.7 | 4.7E-02 | MIF | 0.8 | 1.0E+00 | LYZ | 5.7 | 2.5E-32 | FGFBP2 | 1.1 | 1.0E+00 |
| 20 | DUSP1 | 1.7 | 5.1E-02 | MT-ND6 | 0.8 | 1.0E+00 | RNASE1 | 5.6 | 5.8E-30 | PDCD4 | 1.1 | 1.0E+00 |
| 21 | H1FX | 1.6 | 7.5E-02 | LMNA | 0.8 | 1.0E+00 | FCER1G | 5.5 | 5.0E-29 | HLA-DPB1 | 1.1 | 1.0E+00 |
| 22 | CTSB | 1.6 | 8.2E-02 | HLA-DRB1 | 0.8 | 1.0E+00 | C1orf162 | 5.3 | 3.1E-25 | ZFP36L2 | 1.0 | 1.0E+00 |
| 23 | HLA-DQB1 | 1.6 | 8.9E-02 | JUND | 0.8 | 1.0E+00 | FGL2 | 5.1 | 4.1E-22 | HLA-DPA1 | 1.0 | 1.0E+00 |
| 24 | TNFSF10 | 1.5 | 9.1E-02 | TRBV28 | 0.8 | 1.0E+00 | IFIT2 | 5.0 | 2.3E-21 | DDIT4 | 1.0 | 1.0E+00 |
| 25 | FTL | 1.5 | 9.3E-02 | MT-ND4 | 0.8 | 1.0E+00 | AIF1 | 4.7 | 2.3E-17 | JUNB | 1.0 | 1.0E+00 |

Shared or indicative or clusters 1-2 | Shared or indicative or clusters 3-4 | Shared or indicative or clusters 5-8
