## Extended Data Table 2 for "Multiomic single-cell sequencing defines tissue-specific responses in Stevens-Johnson Syndrome and Toxic epidermal necrolysis"

**Extended Data Table 2. Differential gene signatures of keratinocytes and CD8 Tconv in affected compared to unaffected skin.** The genes that show significant (Wilcoxon rank sum test, Benjamini Hochberg adjusted p <0.05) increased expression are shown in order of fold change. Red font indicates genes significantly differentially expressed but below the threshold for fold change (0.6 log2FC, 1.5 actual fold change). *FDR adj p, False discovery rate adjusted p-value; FC, fold change; Tconv, T conventional cell.*

| | Keratinocyte signature | | | CD8 Tconv signature | | |
|---|---|---|---|---|---|---|
| **Rank** | **Gene** | **FDR adj p** | **log2FC** | **Gene** | **FDR adj p** | **log2FC** |
| 1 | CD74 | 1.2E-34 | 2.0 | GNLY | 2.2E-02 | 1.3 |
| 2 | IFITM3 | 1.2E-22 | 1.8 | CD27 | 1.7E-05 | 0.7 |
| 3 | IFI6 | 6.3E-34 | 1.7 | LY6E | 2.4E-03 | 0.7 |
| 4 | IFITM1 | 2.8E-33 | 1.6 | PSMB9 | 3.6E-04 | 0.6 |
| 5 | HLA-C | 8.6E-12 | 1.3 | RNF213 | 1.4E-03 | 0.6 |
| 6 | HLA-B | 5.2E-05 | 1.1 | LAG3 | 1.5E-03 | 0.6 |
| 7 | B2M | 1.4E-14 | 1.1 | LIMD2 | 9.1E-03 | 0.6 |
| 8 | KRT6B | 4.0E-04 | 1.1 | TIGIT | 1.1E-02 | 0.5 |
| 9 | LY6E | 1.3E-04 | 0.9 | GALNT2 | 4.7E-03 | 0.5 |
| 10 | PSME2 | 1.0E-07 | 0.8 | IFI6 | 1.1E-02 | 0.5 |
| 11 | TMSB10 | 1.3E-04 | 0.8 | LYST | 1.5E-02 | 0.5 |
| 12 | CST3 | 1.3E-04 | 0.8 | XAF1 | 7.2E-04 | 0.4 |
| 13 | DCD | 1.5E-04 | 0.8 | GBP2 | 1.4E-02 | 0.4 |
| 14 | STAT1 | 4.2E-12 | 0.8 | DCD | 4.8E-02 | 0.4 |
| 15 | VIM | 2.1E-04 | 0.7 | ACP5 | 1.1E-02 | 0.4 |
| 16 | GSTP1 | 3.8E-03 | 0.7 | | | |
| 17 | IFI27 | 1.3E-09 | 0.7 | | | |
| 18 | CCL4 | 8.7E-11 | 0.7 | | | |
| 19 | PSME1 | 1.2E-02 | 0.7 | | | |
| 20 | PSMB9 | 2.5E-11 | 0.6 | | | |
| 21 | GAPDH | 8.7E-08 | 0.6 | | | |
