## Extended Data Table 3 for "Multiomic single-cell sequencing defines tissue-specific responses in Stevens-Johnson Syndrome and Toxic epidermal necrolysis"

**Extended Data Table 3. Differential gene signatures of CD8 Tconv expressing dominantly expanded TCR compared to other TCR in affected skin or blister fluid.** The genes that show significant (Wilcoxon rank sum test, Benjamini Hochberg adjusted p <0.05) increased expression above the threshold for fold-change (0.6log2FC) are shown. Genes shared in the signature of dominantly-expanded TCR+ Tconv between affected skin and blister fluid highlighted blue. *TCR, T-cell receptor; FDR adj p, False discovery rate adjusted p-value; FC, fold change; Tconv, T conventional cell.*

| | Dominant TCR in affected skin | | | Dominant TCR in Blister fluid | | |
|---|---|---|---|---|---|---|
| Rank | Gene | FDR adj p | log2FC | Gene | FDR adj p | log2FC |
| 1 | GNLY | 1.0E-06 | 2.7 | GNLY | 1.5E-118 | 1.9 |
| 2 | GZMB | 6.1E-04 | 1.6 | LAG3 | 6.5E-175 | 1.5 |
| 3 | KLRC1 | 3.6E-16 | 1.4 | TIGIT | 6.0E-142 | 1.3 |
| 4 | GALNT2 | 3.3E-07 | 1.4 | KLRC1 | 1.6E-158 | 1.3 |
| 5 | LAG3 | 2.6E-07 | 1.3 | GZMB | 7.7E-65 | 1.1 |
| 6 | BATF | 5.8E-08 | 1.3 | PRF1 | 1.9E-119 | 1.1 |
| 7 | PRF1 | 2.5E-03 | 1.2 | PHLDA1 | 3.3E-150 | 1.0 |
| 8 | DUSP4 | 5.2E-06 | 1.2 | GALNT2 | 6.3E-123 | 1.0 |
| 9 | PTMS | 2.6E-06 | 1.1 | FCGR3A | 8.0E-90 | 1.0 |
| 10 | FCGR3A | 1.2E-03 | 1.0 | ACP5 | 8.9E-100 | 1.0 |
| 11 | TIGIT | 2.3E-03 | 1.0 | NKG7 | 9.1E-102 | 1.0 |
| 12 | ACP5 | 2.8E-06 | 1.0 | HAVCR2 | 1.3E-95 | 0.9 |
| 13 | LYST | 2.1E-05 | 1.0 | IGFLR1 | 4.4E-120 | 0.9 |
| 14 | HAVCR2 | 3.2E-05 | 0.9 | GAPDH | 3.8E-108 | 0.8 |
| 15 | CD27 | 2.1E-02 | 0.9 | ENTPD1 | 2.3E-115 | 0.8 |
| 16 | MT1E | 2.9E-03 | 0.9 | KLRD1 | 1.0E-61 | 0.8 |
| 17 | LAYN | 2.0E-09 | 0.9 | LYST | 3.7E-87 | 0.8 |
| 18 | RAB27A | 2.5E-05 | 0.9 | PTMS | 1.5E-79 | 0.8 |
| 19 | SNX9 | 4.0E-04 | 0.9 | TNFRSF1B | 2.0E-74 | 0.8 |
| 20 | GAPDH | 1.5E-06 | 0.9 | HMOX1 | 2.6E-145 | 0.8 |
| 21 | TNFRSF9 | 9.4E-10 | 0.9 | PKM | 4.7E-63 | 0.8 |
| 22 | CD59 | 8.6E-06 | 0.9 | LAYN | 3.7E-138 | 0.8 |
| 23 | AHI1 | 1.1E-05 | 0.9 | SIRPG | 1.1E-73 | 0.7 |
| 24 | PKM | 1.5E-02 | 0.8 | CCL4 | 5.0E-23 | 0.7 |
| 25 | PHLDA1 | 2.6E-02 | 0.8 | CD27 | 1.4E-66 | 0.7 |
| 26 | CBLB | 1.4E-02 | 0.8 | TNFRSF9 | 8.5E-134 | 0.7 |
| 27 | MTSS1 | 1.2E-08 | 0.8 | CBLB | 7.3E-67 | 0.7 |
| 28 | HMOX1 | 1.2E-09 | 0.8 | CCL3 | 5.7E-68 | 0.7 |
| 29 | ENTPD1 | 2.6E-07 | 0.8 | AD000671.2 | 4.7E-93 | 0.7 |
| 30 | AD000671.2 | 1.2E-03 | 0.8 | ADGRG1 | 1.1E-128 | 0.7 |
| 31 | CTLA4 | 1.8E-05 | 0.8 | CD63 | 3.3E-59 | 0.7 |
| 32 | CD70 | 2.2E-10 | 0.7 | LSP1 | 3.1E-86 | 0.7 |
| 33 | LSP1 | 1.6E-03 | 0.7 | CD8A | 1.5E-42 | 0.7 |
| 34 | PPM1G | 2.0E-02 | 0.7 | S100A4 | 1.3E-50 | 0.7 |
| 35 | APOBEC3C | 4.4E-02 | 0.6 | LINC01943 | 1.6E-82 | 0.7 |
| 36 | CD38 | 2.5E-03 | 0.6 | SERPINB1 | 2.8E-53 | 0.6 |
| 37 | | | | LINC01871 | 2.9E-42 | 0.6 |
| 38 | | | | RHOB | 5.2E-129 | 0.6 |
| 39 | | | | RAB27A | 3.9E-61 | 0.6 |
| 40 | | | | DUSP4 | 7.4E-83 | 0.6 |
| 41 | | | | AC017002.3 | 5.0E-92 | 0.6 |
| 42 | | | | NEAT1 | 5.9E-31 | 0.6 |
| 43 | | | | IFNG | 4.1E-60 | 0.6 |
| 44 | | | | SNX9 | 4.3E-92 | 0.6 |
| 45 | | | | ITGA4 | 1.8E-48 | 0.6 |
| 46 | | | | PGAM1 | 4.0E-48 | 0.6 |
| 47 | | | | CTLA4 | 1.9E-65 | 0.6 |
