## Extended Data Table 4 for "Multiomic single-cell sequencing defines tissue-specific responses in Stevens-Johnson Syndrome and Toxic epidermal necrolysis"

**Extended Data Table 4. Unexpanded clonotypes of CD8+ Tconv cluster 3 with a shared CDRβ sequence with the dominantly-expanded clonotypes.** Unexpanded clonotypes from CD8 Tconv cluster 3 of blister fluids 1-3 are shown that have the exact same CDR3β CASSPDRGGYEQYF sequence as the dominantly-expanded TCR+ population or a single mismatch (underlined). The 'Dual TCR' column indicates if this n=1 count TCR is expressed on a dual TCR+ cell (yes or no). TCRs expressed on the same dual TCR+ cell are indicated by the same symbol (○, ●, ◊, ‡). *Tconv, T conventional cell; TCR, T-cell receptor; TRAV, TCR alpha variable; TRBV, TCR beta variable; TRAJ, TCR alpha joining; TRBJ, TCR beta joining; CDR3, complementary-determining region; Ct., count.*

**Blister fluid 1 (ARM, 338 unexpanded cells from CD8 Tconv cluster 3)**

| TCR | CDR3α | CDR3β | TRAV | TRAJ | TRBV | TRBJ | Ct. | Dual TCR+ |
|---|---|---|---|---|---|---|---|---|
| 1 | CAVQAFRQTGANNLFF | CASSHDRGGYEQYF | TRAV20 | TRAJ36 | TRBV27 | TRBJ2-7 | 1 | yes ○ |
| 2 | CALSEVTTSGTYKYIF | CASSHDRGGYEQYF | TRAV19 | TRAJ40 | TRBV27 | TRBJ2-7 | 1 | yes ○ |
| 3 | CVVATNAGGTSYGKLTF | CASSPDRGGYEQYF | TRAV10 | TRAJ52 | TRBV27 | TRBJ2-7 | 1 | yes |
| 4 | CALSEVTTSGTYKYIF | CASSRDRGGYEQYF | TRAV19 | TRAJ40 | TRBV27 | TRBJ2-7 | 1 | no |
| 5 | CAAIDSWGKLQF | CASSLDRGGYEQYF | TRAV23/DV6 | TRAJ24 | TRBV11-2 | TRBJ2-7 | 1 | yes ‡ |
| 6 | CALSEVRTSGTYKYIF | CASSLDRGGYEQYF | TRAV19 | TRAJ40 | TRBV11-2 | TRBJ2-7 | 1 | yes ‡ |

**Blister fluid 2 (FACE, 181 unexpanded cells from CD8 Tconv cluster 3)**

| TCR | CDR3α | CDR3β | TRAV | TRAJ | TRBV | TRBJ | Ct. | Dual TCR+ |
|---|---|---|---|---|---|---|---|---|
| 1 | CALSEVTTSGTYKYIF | CASSHDRGGYEQYF | TRAV19 | TRAJ40 | TRBV27 | TRBJ2-7 | 1 | yes ○ |
| 2 | CAVQAFRQTGANNLFF | CASSHDRGGYEQYF | TRAV20 | TRAJ36 | TRBV27 | TRBJ2-7 | 1 | yes ○ |
| 3 | CAADTGGFKTIF | CASSPDRGGYEQYF | TRAV13-1 | TRAJ9 | TRBV27 | TRBJ2-7 | 1 | yes ● |
| 4 | CIVRVHSGGGADGLTF | CASSPDRGGYEQYF | TRAV26-1 | TRAJ45 | TRBV27 | TRBJ2-7 | 1 | yes ● |
| 5 | CAVTDNYGQNFVF | CASSPDRGGYEQYF | TRAV25 | TRAJ26 | TRBV27 | TRBJ2-7 | 1 | yes ◊ |
| 6 | CALSANSGNTPLVF | CASSPDRGGYEQYF | TRAV16 | TRAJ29 | TRBV27 | TRBJ2-7 | 1 | yes ◊ |
| 7 | CAVQTNAGNNRKLIW | CASSPDRGGYEQYF | TRAV20 | TRAJ38 | TRBV27 | TRBJ2-7 | 1 | yes |
| 8 | CAVKYTGANSKLTF | CASSPDRGGYEQYF | TRAV12-2 | TRAJ56 | TRBV27 | TRBJ2-7 | 1 | yes |
| 9 | CAAGSSSGTYKYIF | CASSPDRGGYEQYF | TRAV13-1 | TRAJ40 | TRBV27 | TRBJ2-7 | 1 | yes |

**Blister fluid 3 (FOOT, 137 unexpanded cells from CD8 Tconv cluster 3)**

| TCR | CDR3α | CDR3β | TRAV | TRAJ | TRBV | TRBJ | Ct. | Dual TCR+ |
|---|---|---|---|---|---|---|---|---|
| 1 | CALSEVTTSGTYKYIF | CASSHDRGGYEQYF | TRAV19 | TRAJ40 | TRBV27 | TRBJ2-7 | 1 | yes ○ |
| 2 | CAVQAFRQTGANNLFF | CASSHDRGGYEQYF | TRAV20 | TRAJ36 | TRBV27 | TRBJ2-7 | 1 | yes ○ |
| 3 | CAASMTSAGNMLTF | CASSVDRGGYEQYF | TRAV23/DV6 | TRAJ39 | TRBV27 | TRBJ2-7 | 1 | yes □ |
| 4 | CALSEVTTSGTYKYIF | CASSVDRGGYEQYF | TRAV19 | TRAJ40 | TRBV27 | TRBJ2-7 | 1 | yes □ |
| 5 | CALSEVTTSGTYKYIF | CASSYDRGGYEQYF | TRAV19 | TRAJ40 | TRBV27 | TRBJ2-7 | 1 | yes |
| 6 | CALSEVTTSGTYKYIF | CASSFDRGGYEQYF | TRAV19 | TRAJ40 | TRBV27 | TRBJ2-7 | 1 | no |
